# Varstation: a complete and efficient tool to support NGS data analysis

**DOI:** 10.1101/833582

**Authors:** ACO Faria, MP Caraciolo, RM Minillo, TF Almeida, SM Pereira, MC Cervato, JBO Filho

## Abstract

**Summary:** Varstation is a cloud-based NGS data processor and analyzer for human genetic variation. This resource provides a customizable, centralized, safe and clinically validated environment aiming to improve and optimize the flow of NGS analyses and reports related with clinical and research genetics.

**Availability and implementation:** Varstation is freely available at http://varstation.com, for academic use.

**Supplementary information:** <u>Supplementary data</u> are available at *Bioinformatics* online.

## 1 Introduction

In recent years, advances in next-generation sequencing (NGS) technologies and applications, such as whole-genome sequencing (WGS), whole-exome sequencing (WES), and targeted-sequencing have enabled the generation of large-scale data and identification of millions of genetic variants both for research and clinical diagnostic purposes (Yang *et al.*, 2014; Xue *et al.*, 2015). Identifying, interpreting, classifying, and associating human genetics variants with phenotypic particularities among individuals or diseases are important parts of the process to accomplish personalized medicine (Gaspar *et al.*, 2014; Geoffroy *et al.*, 2015).

Data workflow in NGS includes several bioinformatics steps from the analysis of raw sequencing data transforming the signal from the sequencers to raw sequences that are further aligned and compared with the reference genome (Geoffroy *et al.*, 2015). In general, a typical workflow analysis consists of the following steps: raw data quality control (QC), preprocessing, alignment, post-alignment processing, variant calling, annotation, and prioritization (Bao *et al.*, 2014).

There is a pressing need for the development of optimized workflows and variant analysis toolkits that assist clinical and laboratory geneticists and researchers in the identification, classification and ranking of variants in a timely fashion.

We describe a new powerful tool named Varstation (www.varstation.com, Figure 1) to support the human NGS data analysis workflows. Varstation is a cloud-based solution for computational processing and clinical support of NGS genetic testing that provides a customizable, centralized, safe and clinically validated environment. This resource was developed by a multidisciplinary team composed by bioinformaticians, biologists, geneticists, software engineers, and designers. The goal was to design a resource that could be used for different analysis workflows being both scientifically sound and easy to use.

**Figure 1:**
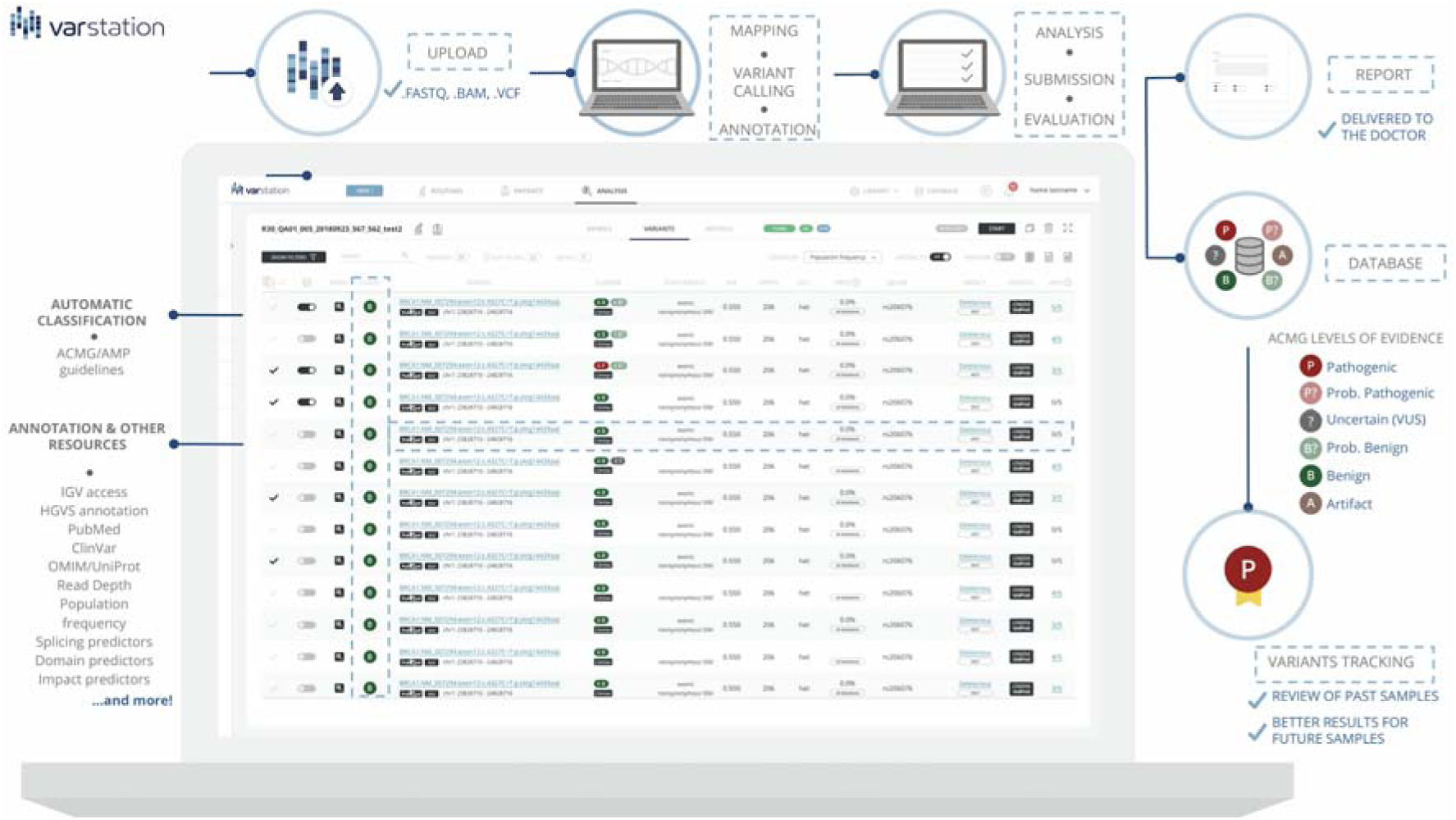
Layout and main features of Varstation.

## 2 Software description

### 2.2 The Varstation

Varstation comprises three main components of bioinformatics pipelines: alignment of the sequenced readings to the reference genome; variant calling and annotation, which entails capturing and summarizing all existing information about the variant across multiple public databases.

The strength of Varstation workflow is based on (i) flexibility: files from any DNA sequencing platform, in any format (.fastq, .bam, and .vcf) are processed to generate results; (ii) end-to-end and automated processing: evaluation of quality parameters, mapping, multiple variant callers, database annotation and automatic variant pre-classification according recommendations and guidelines of the American College of Medical Genetics and Genomics (ACMG), the Association of Molecular Pathology (AMP), the College of American Pathologists (CAP) (Richards *et al.*, 2015) and the Clinical Laboratory Accreditation Program (*Programa de Acreditação de Laboratórios Clínicos*, PALC) (http://www.sbpc.org.br/programas-da-qualidade/documentos-do-palc/); (iii) support for clinical interpretation: more than 100 genetic mutations databases are incorporated, including data from germline, somatic and structural variants; (iv) robust filters: filtering engine based on all annotated mutation data; (v) clear and structured results: relevant clinical information to support the medical report. In addition, Varstation provides visualization of data features to share results with other institutions. Information from the external databases and filters provides in Varstation are described in <u>supplementary table S1.</u>

### 2.3 Workflow optimization: main features

To access Varstation, the user must register using a valid e-mail address. After an e-mail confirmation, the new user will be able to access the platform and the new users onboarding tutorial (raw clinical exome data sample are available). Users can update and personalize the workflow depending on the type of the starting file and adjust parameters each component.

QC of raw data and their preprocessing can be performed using the FastQC (Wingett and Andrews, 2018), BEDTools (Quinlan and Hall, 2010), BamTools (Barnett *et al.*, 2011) and VCFtools (Danecek *et al.*, 2011) toolkits. After raw data QC and preprocessing, the next step is to map readings to the reference genome and process the variant calling with high efficiency and accuracy. Two of the main mapping and alignment tools are available in the platform: Burrows–Wheeler Aligner (BWA) (Li, 2013) and Torrent Mapping Alignment Program (TMAP, https://github.com/iontorrent/TS/tree/master/Analysis/TMAP).

Programs available for germline variant calling include GATK - UnifiedGenotyper and HaplotypeCaller (Van der Auwera *et al.*, 2013), SAMtools (Li *et al.*, 2009), FreeBayes (Garrison and Marth, 2012), Atlas (Challis *et al.*, 2012) and smCounter (Xu *et al.*, 2018). For somatic variant detection, Varstation provides GATK (Van der Auwera *et al.*, 2013), SAMtools (Li *et al.*, 2009), VarScan (Koboldt *et al.*, 2012), Vardict (Lai *et al.*, 2016) and smCounter (Xu *et al.*, 2018).

After variant identification, the annotation attributes such as genomic components, gene symbol, amino acid change, and functional consequences are attached to the variant list. For annotation, Varstation offers one in-house annotation algorithm that was based on ANNOVAR, the most commonly used variant annotation programs (Wang *et al.*, 2010).

The last step is variant filtration and prioritization. Users can filter the variants by gene name, ACMG (Richards *et al.*, 2015) or ClinVar (Landrum *et al.*, 2018) classification, protein coding consequence, variant allele frequency (VAF), depth coverage, frequency in public databases, specifically: 1000g, ESP6500, gnomAD, ABraOM, and ExAC (1000 Genomes Project Consortium *et al.*, 2010; Tennessen *et al.*, 2012; Karczewski *et al.*, 2019; Naslavsky *et al.*, 2017; Lek *et al.*, 2016), presence in dbNSP (Sherry *et al.*, 2001), protein-damage prediction tools (VEST, FATHMM, SIFT, Polyphen, PROVEAN, CADD) (Douville *et al.*, 2016; Carter *et al.*, 2013; Shihab *et al.*, 2014; Shihab, Gough, Cooper, Day, *et al.*, 2013; Shihab, Gough, Cooper, Stenson, *et al.*, 2013; Sim *et al.*, 2012; Adzhubei *et al.*, 2013; Choi and Chan, 2015; Rentzsch *et al.*, 2019), disease associated by OMIM (McKusick, 2007) and UniProt (UniProt Consortium, 2019). Variants are also linked to any associated phenotypes in the Human Phenotype Ontology (HPO) (Köhler *et al.*, 2019). Users can also easily visualize the variant of interest, in their context of the .bam files, using the Integrative Genomics Viewer (Robinson *et al.*, 2017).

One of the key features of Varstation is that users can build their organization’s internal database, customizing their own filter’s pipelines, link variants to phenotypes, diseases or articles, and can make their own pathogenicity assessments, potentially allowing for the rapid identification of recurrent mutations, and periodic variant reevaluation or reanalysis, following the ACMG guidelines (Deignan *et al.*, 2019). This feature is extremely relevant mainly for clinical laboratories that need policies and protocols established for variant re-analyzes. Besides that, Varstation also provides multiple ready-to-use standardized pipelines.

Finally, Varstation enables the analysis of each sample by up to 3 experts to guarantee accuracy of results, and then it is possible to generate a customized report with a list of candidate variants that will be reported.

## 4 Conclusion

The process of analyzing NGS data in Varstation improves the performance, traceability, and safety of analyses, in addition the resource complies with the good international practices for analysis of human genetic variants.

Varstation is a complete, simple, and powerful tool to process and analyze NGS data that was already adopted by leading laboratories in Brazil. The initial project of Varstation was developed in June 2015 and became available for use only for the *Sociedade Beneficiente Hospital Israelita Albert Einstein* hospital and the *Genomika* laboratory. In 2018, Varstation began to become available to other laboratories, hospitals, and research centers. Since then, Varstation has being used by approximately 500 organizations and has processed more than 10,000 samples.

As one of Varstation best features, the software has an online chat service that helps users to solve problems found during the analysis process. In addition, a multi-disciplinary team performs hands-on courses to help the community to further improve the quality of their genomic analysis.

## Acknowledgments

We would like to thank all colleagues from the Clinical Laboratory of Hospital Israelita Albert Einstein for the partnership and the enriching discussions. We would like to also thank to all developers from the Innovation Laboratory that did contribute and keep working in Varstation, offering an incredible atmosphere of work.

