## Supplemental Table 1 for "Varstation: a complete and efficient tool to support NGS data analysis"

| Filter type | Filter Name | Source | URL |
| --- | --- | --- | --- |
| Public Genomic Databases |  |  |  |
|  | 1000Genomes | [IGSR: The International Genome Sample Resource](http://www.internationalgenome.org/) | <http://www.internationalgenome.org/1000-genomes-browsers> |
|  | ABRaOM | Human Genome and Stem Cell Research Center - HUG-CELL | <http://abraom.ib.usp.br/> |
|  | ESP6500 | NHLBI GO Exome Sequencing Project (ESP) | <https://esp.gs.washington.edu/drupal/> |
|  | ExAC | Exome Aggregation Consortium | <http://exac.broadinstitute.org/> |
|  | gnomAD | Genome Aggregation Database | <https://gnomad.broadinstitute.org/> |
|  | HRC | [The Haplotype Reference Consortium](http://www.haplotype-reference-consortium.org/) | <http://www.haplotype-reference-consortium.org/> |
|  | ICGC | the International Cancer Genome Consortium | <https://icgc.org/> |
|  | Known VARiants | KnownVar | <https://redmine.igm.cumc.columbia.edu/projects/atav/wiki/KnownVar> |
| General filters |  |  |  |
|  | Allele Frequency |  |  |
|  | Allele Frequency (UMI) |  |  |
|  | Alternative Forward |  |  |
|  | Alternative Reverse |  |  |
|  | Coding Change |  |  |
|  | genomicSuperDups | UCSC | <http://varianttools.sourceforge.net/Annotation/GenomicSuperDups> |
|  | Interpro_domain | InterPro: protein sequence analysis & classification | <https://www.ebi.ac.uk/interpro/> |
|  | LRG | Locus Reference Genomic | <https://www.lrg-sequence.org/> |
|  | Protein Change |  |  |
|  | Read Depth |  |  |
|  | Read Depth (UMI) |  |  |
|  | Reference Forward |  |  |
|  | Reference Reverse |  |  |
|  | RepeatMasker | RepeatMasker Web Server | <http://www.repeatmasker.org/> |
|  | Strand Bias |  |  |
|  | Var Call Count |  |  |
|  | Zygosity |  |  |
| Variant consequence |  |  |  |
|  | Alternative Allele |  |  |
|  | Amino Acid Change |  |  |
|  | Chromossome |  |  |
|  | DbSNP | Single Nucleotide Polymorphism database - NCBI | <https://www.ncbi.nlm.nih.gov/projects/SNP/> |
|  | End |  |  |
|  | Exonic Functional Consequence |  |  |
|  | Functional Category |  |  |
|  | Gene Detail |  |  |
|  | Gene Name |  |  |
|  | HGVS | [Human Genome Variation Society](http://www.hgvs.org/) | <http://www.hgvs.org/mutnomen/> |
|  | PFAM | EMBL-EBI | <https://pfam.xfam.org/> |
|  | Reference |  |  |
|  | snp138 | Single Nucleotide Polymorphism database - NCBI | <https://www.ncbi.nlm.nih.gov/projects/SNP/> |
|  | Start |  |  |
|  | wgEncodeBroadHmmGm12878HMM | Encode/NHGRI | <https://www.encodeproject.org/> |
| Protein |  |  |  |
|  | CADD_phred | Combined Annotation Dependent Depletion | <https://cadd.gs.washington.edu/info> |
|  | CADD_raw | Combined Annotation Dependent Depletion | <https://cadd.gs.washington.edu/info> |
|  | CADD_raw_rankscore | Combined Annotation Dependent Depletion | <https://cadd.gs.washington.edu/info> |
|  | DANN_rankscore | deleterious annotation of genetic variants using neural networks | <https://cbcl.ics.uci.edu/public_data/DANN/> |
|  | DANN_score | deleterious annotation of genetic variants using neural networks | <https://cbcl.ics.uci.edu/public_data/DANN/> |
|  | Eigen_coding_or_noncoding | Columbia university | <http://www.columbia.edu/~ii2135/eigen.html> |
|  | Eigen-PC-raw | Columbia university | <http://www.columbia.edu/~ii2135/eigen.html> |
|  | Eigen-raw | Columbia university | <http://www.columbia.edu/~ii2135/eigen.html> |
|  | FATHMM_converted_rankscore | Functional Analysis through Hidden Markov Models/University of Bristol | <http://fathmm.biocompute.org.uk/> |
|  | fathmm-MKL_coding_pred | Functional Analysis through Hidden Markov Models/University of Bristol | <http://fathmm.biocompute.org.uk/> |
|  | fathmm-MKL_coding_rankscore | Functional Analysis through Hidden Markov Models/University of Bristol | <http://fathmm.biocompute.org.uk/> |
|  | fathmm-MKL_coding_score | Functional Analysis through Hidden Markov Models/University of Bristol | <http://fathmm.biocompute.org.uk/> |
|  | FATHMM_pred | Functional Analysis through Hidden Markov Models/University of Bristol | <http://fathmm.biocompute.org.uk/> |
|  | FATHMM_score | Functional Analysis through Hidden Markov Models/University of Bristol | <http://fathmm.biocompute.org.uk/> |
|  | GenoCanyon_score | GenoCanyon | <http://genocanyon.med.yale.edu/> |
|  | GenoCanyon_score_rankscore | GenoCanyon | <http://genocanyon.med.yale.edu/> |
|  | integrated_confidence_value | FitCons/Abriza  Cold Spring Harbor Lab | <http://compgen.cshl.edu/fitCons/> |
|  | integrated_fitCons_score | FitCons/Abriza  Cold Spring Harbor Lab | <http://compgen.cshl.edu/fitCons/> |
|  | integrated_fitCons_score_rankscore | FitCons/Abriza  Cold Spring Harbor Lab | <http://compgen.cshl.edu/fitCons/> |
|  | LRT_converted_rankscore | Likelihood ratio test | <http://evomics.org/resources/likelihood-ratio-test/> |
|  | LRT_pred | Likelihood ratio test | <http://evomics.org/resources/likelihood-ratio-test/> |
|  | LRT_score | Likelihood ratio test | <http://evomics.org/resources/likelihood-ratio-test/> |
|  | M-CAP_pred | M-CAP | doi: 10.1038/ng.3703 |
|  | M-CAP_rankscore | M-CAP | doi: 10.1038/ng.3703 |
|  | M-CAP_score | M-CAP | doi: 10.1038/ng.3703 |
|  | MetaLR_pred | MetaLR/Coco Dong  USC Biostatiscs Department | <http://wglab.org/members/15-member-detail/36-coco-dong> |
|  | MetaLR_rankscore | MetaLR/Coco Dong  USC Biostatiscs Department | <http://wglab.org/members/15-member-detail/36-coco-dong> |
|  | MetaLR_score | MetaLR/Coco Dong  USC Biostatiscs Department | <http://wglab.org/members/15-member-detail/36-coco-dong> |
|  | MetaSVM_pred | MetaSVM/Coco Dong  USC Biostatiscs Department | <http://wglab.org/members/15-member-detail/36-coco-dong> |
|  | MetaSVM_rankscore | MetaSVM/Coco Dong  USC Biostatiscs Department | <http://wglab.org/members/15-member-detail/36-coco-dong> |
|  | MetaSVM_score | MetaSVM/Coco Dong  USC Biostatiscs Department | <http://wglab.org/members/15-member-detail/36-coco-dong> |
|  | MutationAssessor_pred | MutationAssessor | <http://mutationassessor.org/r3/> |
|  | MutationAssessor_score | MutationAssessor | <http://mutationassessor.org/r3/> |
|  | MutationAssessor_score_rankscore | MutationAssessor | <http://mutationassessor.org/r3/> |
|  | MutationTaster_converted_rankscore | MutationTaster | <http://www.mutationtaster.org/> |
|  | MutationTaster_pred | MutationTaster | <http://www.mutationtaster.org/> |
|  | MutationTaster_score | MutationTaster | <http://www.mutationtaster.org/> |
|  | Polyphen2_HDIV_pred | Polymorphism Phenotyping v2 HDIV | <http://genetics.bwh.harvard.edu/pph2/> |
|  | Polyphen2_HDIV_rankscore | Polymorphism Phenotyping v2 HDIV | <http://genetics.bwh.harvard.edu/pph2/> |
|  | Polyphen2_HDIV_score | Polymorphism Phenotyping v2 HDIV | <http://genetics.bwh.harvard.edu/pph2/> |
|  | Polyphen2_HVAR_pred | Polymorphism Phenotyping v2 HVAR | <http://genetics.bwh.harvard.edu/pph2/> |
|  | Polyphen2_HVAR_rankscore | Polymorphism Phenotyping v2 HVAR | <http://genetics.bwh.harvard.edu/pph2/> |
|  | Polyphen2_HVAR_score | Polymorphism Phenotyping v2 HVAR | <http://genetics.bwh.harvard.edu/pph2/> |
|  | PROVEAN_converted_rankscore | Protein Variation Effect Analyzer)/J. Craig Venter Intitute | <http://provean.jcvi.org/index.php> |
|  | PROVEAN_pred | Protein Variation Effect Analyzer)/J. Craig Venter Intitute | <http://provean.jcvi.org/index.php> |
|  | PROVEAN_score | Protein Variation Effect Analyzer)/J. Craig Venter Intitute | <http://provean.jcvi.org/index.php> |
|  | SIFT_converted_rankscore | Sorting intolerated from tolerated/Hutchinson  Cancer Research Center | <https://sift.bii.a-star.edu.sg/> |
|  | SIFT_pred | Sorting intolerated from tolerated/Hutchinson  Cancer Research Center | <https://sift.bii.a-star.edu.sg/> |
|  | SIFT_score | Sorting intolerated from tolerated/Hutchinson  Cancer Research Center | <https://sift.bii.a-star.edu.sg/> |
|  | VEST3_rankscore | Variant Effect Scoring Tool/John Hopkins University | <https://karchinlab.org/apps/appVest.html> |
|  | VEST3_score | Variant Effect Scoring Tool/John Hopkins University | <https://karchinlab.org/apps/appVest.html> |
| Disease associated |  |  |  |
|  | CliVar Accession | ClinVar NCBI | <https://www.ncbi.nlm.nih.gov/clinvar/> |
|  | ClinVar Database | ClinVar NCBI | <https://www.ncbi.nlm.nih.gov/clinvar/> |
|  | ClinVar Disease | ClinVar NCBI | <https://www.ncbi.nlm.nih.gov/clinvar/> |
|  | ClinVar Disease ID | ClinVar NCBI | <https://www.ncbi.nlm.nih.gov/clinvar/> |
|  | ClinVar Significance | ClinVar NCBI | <https://www.ncbi.nlm.nih.gov/clinvar/> |
|  | COSMIC | Catalogue Of Somatic Mutations In Cancer | <https://cancer.sanger.ac.uk/cosmic> |
|  | GTEx_V6_gene | Genotype-Tissue Expression project | <https://www.gtexportal.org/home/index.html> |
|  | GTEx_V6_tissue | Genotype-Tissue Expression project | <https://www.gtexportal.org/home/index.html> |
|  | InterVar(automated) | InterVar | <http://wintervar.wglab.org/> |
|  | OMIM_genemap2 | OMIM | [https://www.omim.org](https://www.omim.org/) |
|  | regsnp_disease | regSNPs: a strategy for prioritizing regulatory single nucleotide substitutions | <https://academic.oup.com/bioinformatics/article/28/14/1879/218705> |
|  | regsnp_fpr | regSNPs: a strategy for prioritizing regulatory single nucleotide substitutions | <https://academic.oup.com/bioinformatics/article/28/14/1879/218705> |
|  | UNIPROT | UniProt | <https://www.uniprot.org/> |
| Splicing consequence |  |  |  |
|  | dbscSNV_ADA_SCORE | Database of Non-Synonymous Functional Polymorphisms | <https://sites.google.com/site/jpopgen/dbNSFP> |
|  | dbscSNV_RF_SCORE | Database of Non-Synonymous Functional Polymorphisms | <https://sites.google.com/site/jpopgen/dbNSFP> |
|  | regsnp_splicing_site | regSNPs: a strategy for prioritizing regulatory single nucleotide substitutions | <https://academic.oup.com/bioinformatics/article/28/14/1879/218705> |
|  | Spidex Dpsi_max_tissue | SPIDEX | <https://www.deepgenomics.com/spidex/> |
|  | Spidex Dpsi_zscore | SPIDEX | <https://www.deepgenomics.com/spidex/> |
| Nucleotide conservation |  |  |  |
|  | GERP++_RS | Genomic Evolutionary Rate Profiling/Starford University | <http://mendel.stanford.edu/SidowLab/downloads/gerp/index.html> |
|  | GERP++_RS_rankscore | Genomic Evolutionary Rate Profiling/Starford University | <http://mendel.stanford.edu/SidowLab/downloads/gerp/index.html> |
|  | phastCons100way_vertebrate_rankscore | Phylogenetic Analysis with Space/Time Models | <http://compgen.cshl.edu/phast/> |
|  | phastCons20way_Mammalian | Phylogenetic Analysis with Space/Time Models | <http://compgen.cshl.edu/phast/> |
|  | phastCons20way_mammalian_rankscore | Phylogenetic Analysis with Space/Time Models | <http://compgen.cshl.edu/phast/> |
|  | phastCons7way_Vertebrate | Phylogenetic Analysis with Space/Time Models | <http://compgen.cshl.edu/phast/> |
|  | phyloP100way_vertebrate_rankscore | PhyloP basewise conservation | <https://ccg.epfl.ch/mga/hg19/phylop/phylop.html> |
|  | phyloP20way_Mammalian | PhyloP basewise conservation | <https://ccg.epfl.ch/mga/hg19/phylop/phylop.html> |
|  | phlyloP20way_Mammalian_rankscore | PhyloP basewise conservation | <https://ccg.epfl.ch/mga/hg19/phylop/phylop.html> |
|  | phyloP7way_vertebrate | PhyloP basewise conservation | <https://ccg.epfl.ch/mga/hg19/phylop/phylop.html> |
|  | SiPhy29way_logOdds | 29 Mammals Project - Broad Institute | <https://www.broadinstitute.org/mammals-models/29-mammals-project-supplementary-info> |
|  | SiPhy29way_logOdds_rankscore | 29 Mammals Project - Broad Institute | <https://www.broadinstitute.org/mammals-models/29-mammals-project-supplementary-info> |
| Vars-Data Filters |  |  |  |
|  | Variant Allele Frequency maximum in the pipeline |  |  |
|  | Variant Allele Frequency in the routine |  |  |
|  | Variant Allele Frequency medium in the pipeline |  |  |
|  | Variant Allele Frequency medium in the routine |  |  |
|  | Variant Allele Frequency median in the pipeline |  |  |
|  | Variant Allele Frequency median in the routine |  |  |
|  | Variant Allele Frequency minimum in the pipeline |  |  |
|  | Variant Allele Frequency minimum in the routine |  |  |
|  | Routine samples |  |  |
|  | Pipeline samples |  |  |
|  | Routine standard deviation |  |  |
|  | Variant frequency - pipeline |  |  |
|  | Variant frequency - routine |  |  |
|  | Pipeline variant |  |  |
|  | Routine variant |  |  |
|  | Routine variant (heterozygous) |  |  |
|  | Routine variant (homozygous) |  |  |
